# Using Environmental DNA to Reconstruct Amphibian Communities at Sites Infected with *Batrachochytrium salamandrivorans* in the Netherlands

**DOI:** 10.1101/2023.11.22.568296

**Authors:** Anna M. Davison, Annemarieke Spitzen–van der Sluijs, Matthew J. O’Donnell, Andhika P. Prasetyo, Holly A. Broadhurst, Naiara G. Sales, Jelger E. Herder, Ilaria Coscia, Allan D. McDevitt

## Abstract

The recently discovered *Batrachochytrium salamandrivorans* (*Bsal*) fungus can cause high mortality rates in some European salamanders and newts (urodelans) and has the potential to expand its currently small invasive range in Europe. Therefore, monitoring its distribution and better understanding both the species threatened and the mechanics of infection are essential in mitigating the damage *Bsal* may cause. Environmental DNA (eDNA) has emerged as a promising non-invasive method for detecting both this fungal pathogen and the amphibian communities in infected areas. We applied these methods in the province Gelderland, the Netherlands where the pathogen has previously been detected and is expanding its range, with the goal of detecting the natural amphibian community present. We sampled 27 water bodies in the region surrounding the known outbreak sites. We used data from a *Bsal*-specific qPCR assay to determine its presence-absence and applied an eDNA metabarcoding approach to characterize the amphibian communities using two different primer sets. The 12S vertebrate primer set outperformed the 16S amphibian primer set and detected all the expected amphibians in the study area: *Bufo bufo, Lissotriton vulgaris, Pelobates fuscus, Pelophylax spp*., *Rana temporaria* and *Triturus cristatus*. *Bsal* was detected at eight of the ponds. A distance-based redundancy analysis found a weak but significant relationship between *Bsal* presence and the composition of amphibian communities using eDNA. This study provides the foundation for future studies on *Bsal* and its relationship with amphibian communities in Europe, highlighting the need for further research into the mechanisms of persistence and transmission between water bodies.

## Introduction

Amphibians are the most threatened vertebrate class with 40.7% of species classified as threatened according to the International Union for Conservation of Nature’s (IUCN) Red List of Threatened Species (Luedtke et al., 2023). Among the causes of this decline are deforestation, habitat loss and climate change alongside pathogens like ranavirus and the virulent fungal disease chytridiomycosis. *Batrachochytrium dendrobatidis* (*Bd*) was, until recently, thought to be the only fungal disease causing chytridiomycosis. In 2010, a dramatic mortality event of fire salamanders (*Salamandra salamandra*) started in the Netherlands and left just 0.01% of the Dutch population remaining by 2016 (Spitzen-van der Sluijs et al., 2016). The superficial skin lesions and deep epidermal ulcerations found on these salamanders were caused by a second highly pathogenic chytrid fungus, *Batrachochytrium salamandrivorans* (*Bsal*) (Martel et al., 2013). As the fungus breaks down the skin bacteria, it can cause septicemia and death (Bletz et al., 2018). The Dutch NGO RAVON (Reptile, Amphibian and Fish Conservation the Netherlands) has been heavily involved in the monitoring of *Bsal* in the Netherlands since its discovery. Since 2010, the fungus has spread within the Netherlands to four new locations outside of the original infection site in Limburg. Thought to have been introduced from Asia to Europe via the pet trade, *Bsal* has now been found in captive populations in five European countries of which Germany, the Netherlands and Belgium also host wild populations of infected urodelans (Martel et al., 2014; Spitzen-van der Sluijs et al., 2016; Laking et al., 2017; EFSA Panel on Animal Health and Welfare, 2018). Its known range was expanded in 2020 to include Spain after a mass mortality of marbled newts (*Triturus marmoratus*) was attributed to *Bsal* infection (Martel et al., 2020).

Although *Bsal* and *Bd* are superficially similar in their cause of lethal skin erosion, studies show that potential urodelan hosts show different reactions to each fungus when exposed (Martel et al., 2013, 2014). Furthermore, there is a deficit in the knowledge of the pathogenic mechanisms of *Bsal* and the influence of biotic and abiotic factors on pathogen transmission is largely undocumented (Blaustein et al., 2018). The current scientific opinion on *Bsal* transmission detailed in a paper from the EFSA Panel on Animal Health and Welfare (2018) suggests that the most important vectors of transmission within a subpopulation are via active and passive amphibian carriers. Several species of anuran have been shown to carry the fungus in the captive pet trade (Nguyen et al., 2017) and experimentally (Stegen et al.,, 2017), but currently only *Rana temporaria* has been shown to do so in the wild outside the native range of *Bsal* (EFSA et al.,, 2018; Lötters et al., 2020). More tolerant urodelans can also act as vectors and reservoirs for the fungus, such as the Alpine newt (*Ichthyosaura alpestris*) which could be exacerbating the decline of susceptible local *S. salamandra* populations according to a recent study (Beninde et al., 2021). However, transfer via amphibian-related human activities and wild water birds have the most potential when considering larger distances of transfer between metapopulations (Zhu et al., 2014; EFSA Panel on Animal Health and Welfare, 2018). Experimental evidence has shown that *Bsal* can adhere to the legs of geese to transfer larger distances between ponds (Stegen et al., 2017).

*Bsal* is likely to be more widespread through Europe than is currently known as there is limited surveillance of the disease. Although individuals can be tested reliably using skin swabs and qPCR (quantitative or real-time PCR), this is only generally done after a decline in a population is recorded or dead individuals are found (Blooi et al., 2013). Current active and passive surveillance techniques are not efficient as *Bsal* has a widespread and scattered distribution, can be present in low concentrations, and sick or dead individuals can be difficult to find (Spitzen-van der Sluijs et al., 2016; 2018; Dalbeck et al., 2018). However, a recent paper from Spitzen-van der Sluijs et al. (2020) detailed a new technique utilizing environmental DNA (eDNA) to overcome many of the issues associated with monitoring *Bsal*. eDNA is DNA shed from organisms as they interact with the environment and can be found in samples of water, soil and air (Roh et al., 2006; Ficetola et al., 2008; Lynggaard et al., 2022). This eDNA can then be analyzed using a variety of new and emerging techniques, broadly categorized into single-species or metabarcoding approaches, to determine the species present at a sample’s site of origin. The single-species approach is also known as a species-specific or targeted approach and is used to determine the presence of eDNA from a single species of interest within a sample using species-specific primers and probes (Thomsen et al., 2012). However, the potential for eDNA to inform research on *Bsal* is not limited to single-species monitoring of the fungus itself. eDNA metabarcoding has the capacity to detect full communities of species and can therefore be utilized to determine the community of amphibians present in a *Bsal* infected area (Charvoz et al., 2021; Svenningsen et al., 2022). Attaining information on the full amphibian community present at sites of *Bsal* infection using metabarcoding is beneficial for a number of reasons including both identifying the species at risk and the potential vector species. Furthermore, through establishment of a chronosequence or multiple sampling events over time, changes to the amphibian community composition as a result of *Bsal* and *Bsal*-caused population declines can be identified.

In this study, we aim to use eDNA metabarcoding to characterize the amphibian communities present at multiple sites in the Netherlands where *Bsal* is both present and absent in close proximity to one another. We then determine whether the composition of these communities is associated with the presence of *Bsal* at a fine spatial scale to investigate possible means of transmission and infection.

## Materials and Methods

Two ponds were identified which have been confirmed to contain *Bsal* (RAVON, 2021). Twenty-seven water bodies within a 2 km radius of the original two sites were subject to one session of eDNA sampling in mid-April 2021. This did not include the original two water bodies. Figure 1 shows the location of the study area within the Netherlands, but the exact location is anonymized to obscure the location of protected species and because many of the sites are on private property.

**Fig. 1.**
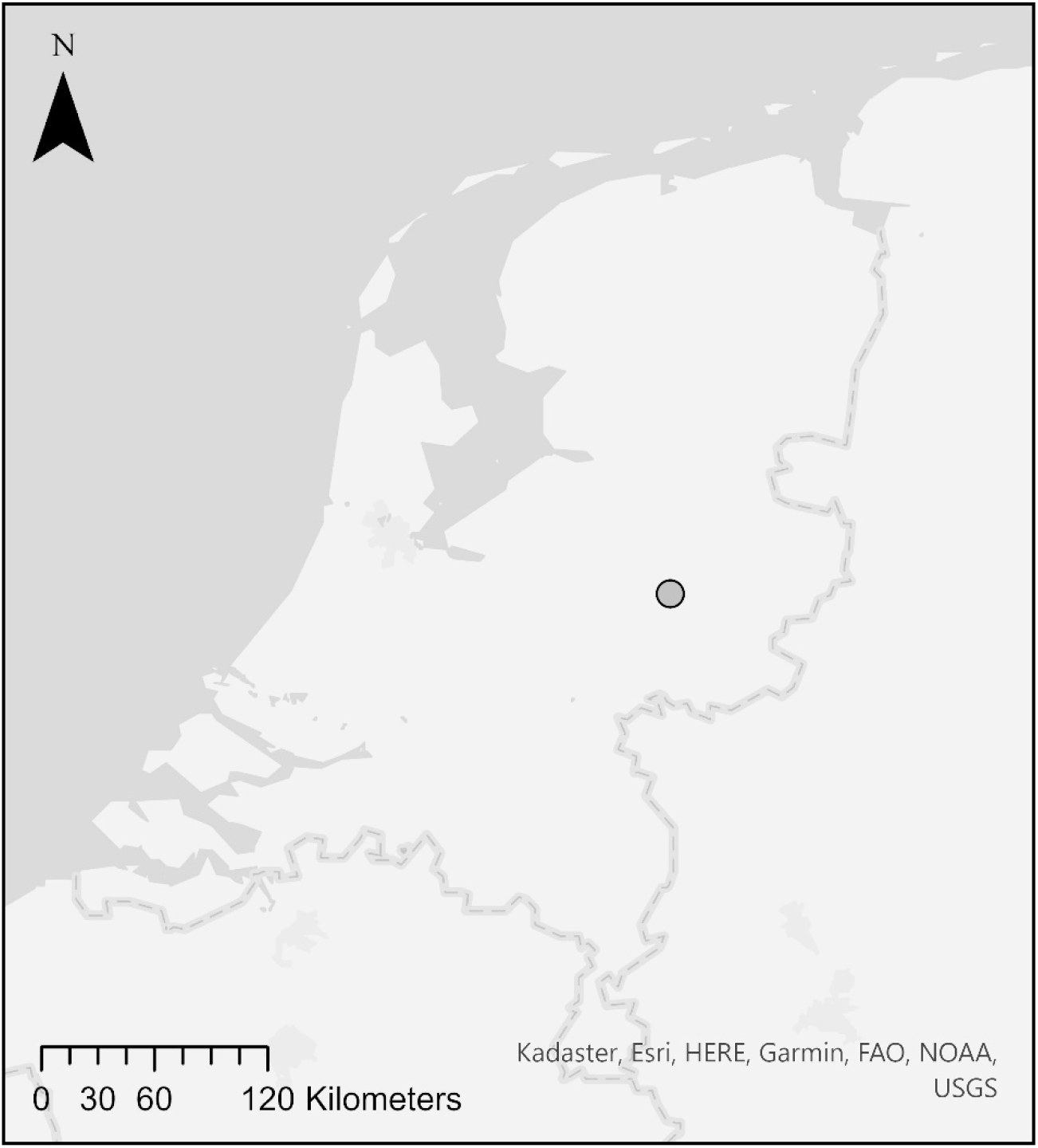
Approximate location of the study site within the Netherlands

For each pond, 20 systematic samples of 100 mL of water were taken standing on the banks of the pond using a 100 mL sterile water sampling ladle as per Spitzen-van der Sluijs et al. (2020). In brief, these water samples were collected from the top 5 cm of the water, decanted into a 2 L self-supporting sterile Whirl-Pak® bag, shaken to homogenize the liquid and the full 2 L of water was filtered through a VigiDNA® 0.45 µM cross-flow filtration capsule. The filtration capsule was then filled with 80 mL of CL1 Conservation buffer to preserve the filter. Samples were then stored at room temperature and sent from RAVON to SPYGEN for single-species eDNA analysis detecting *Bsal*. Full details of the DNA extraction methods and single-species qPCR assay to detect *Bsal* are given in Spitzen-van der Sluijs et al. (2020). The remaining extracted DNA was then sent on to the University of Salford, UK on dry ice for the following eDNA metabarcoding analyses.

We amplified the extracted DNA samples on two separate occasions using two different metabarcoding primer sets. The first was the vertebrate 12S-V5 primer set (12S-V5_F 5’ TAGAACAGGCTCCTCTAG 3’; 12S-V5_R 5’ TTAGATACCCCACTATGC 3’; Riaz et al., 2011) which amplifies a short fragment (∼98 bp) of the 12S rRNA region with sample specific multiplex identifier (MIDs) tags. During PCR for these primers, the samples were denatured for 5 min at 95 °C followed by 35 cycles of 15 s at 95 °C, 30 s at 57 °C and 30 s at 72 °C with a final elongation of 5 min at 72°. The second was the 16S amphibian primer set which targets a 150 bp fragment in the 16S rRNA region of the mitochondrial genome (BA-4445-F 5’-RACCGTGCRAAGGTAGCR-3; BA-178-R 5’-CCATRGGGTCYTCTCGTCT-3’; Bálint et al., 2018) with sample specific MIDs tags. The PCR amplification for these primers followed an adapted protocol with samples denatured for 15 min at 95 °C followed by 35 cycles of 30 s at 94°C, 1 min 30 s at 55 °C and 1 min at 72 °C with a final elongation of 30 min at 60 °C. Hereafter, these primer sets will be referred to 12S vertebrate and 16S amphibian throughout. Twenty-four unique MIDs tags were used for each primer set, for both forward and reverse primers, to differentiate between samples. During PCR for both primer sets, each sample had three replicates to reduce biases in individual reactions. Three replicates of a positive control sample from another project involving captive *Agalychnis lemur*, a Central American species, were included on the plate for each primer set to identify tag jumping for subsequent data filtering steps alongside four negative control replicates to identify potential cross-sample contamination during the PCR process. Amplification was confirmed using 1.2% agarose gel electrophoresis stained with GelRed (Cambridge Bioscience). PCR products were then pooled into two separate libraries, one for each primer set. Full details of the library preparation, sequencing and bioinformatics can be found in the Supplementary Material.

All statistical analyses were conducted in R v4.1.2 (R Core Team, 2021) and maps were made in ArcGIS Pro 2.9 (ESRI, 2021) and Adobe Illustrator 26.0 (Adobe, 2021). Maps of the amphibian communities detected at each site by the two primer sets were constructed to show their distribution alongside the presence of *Bsal*. Subsequent analyses were conducted with the data converted into presence-absence and site 22 was removed as no data for this site was attained, likely due to an error at the PCR stage. A species accumulation curve was constructed using the iNEXT function (Hsieh et al., 2016), extrapolating the amphibian species richness data to 50 sites for each primer set and the combined primer data to determine their effectiveness.

Sites were split into those which tested positive and those which tested negative for *Bsal*. Sites with no species detected were removed (sites 2, 16, 17 and 19) which left 15 *Bsal* negative sites and 7 *Bsal* positive sites. The assumption of no difference in dispersion between *Bsal* positive and negative amphibian communities was tested using a permutational analysis of multivariate dispersions (PERMDISP) with 999 permutations and a significant difference was found (F = 10.44, p = 0.005) (Oksanen et al., 2019). As PERMANOVA is not as powerful with a significant PERMDISP result, a distance-based redundancy analysis (dbRDA) was performed instead, using the Jaccard index matrix as the response variable. This was done to determine the strength of *Bsal* as an explanatory variable of amphibian community composition. The significance of the results was assessed with an analysis of variance (function ANOVA in vegan) with 1,000 permutations.

## Results

The *Bsal* qPCR assay identified 8 *Bsal* positive sites and 19 negative sites. The Miseq sequencing runs yielded ∼1.7 million raw reads for the 12S vertebrate primers and ∼1 million reads for the 16S amphibian primers prior to data filtering. Table S1 shows the reads for each primer dataset remaining at the end of the bioinformatics process and after each stage of the filtering process. Non-target reads (non-vertebrate) comprised just 1.3% for the 12S vertebrate primer data, with 498,302 reads belonging to the class Amphibia (Table S1). See the Supplementary Material and Table S2 for a breakdown on the vertebrate communities found at each site. The vast majority of reads sequenced in the 16S amphibian primer dataset were not amphibian as shown by the dramatic reduction in reads at the final filtering step (Table S1). The reads for the 16S amphibian primers were dominated by Rotifera (27.4%), Arthropoda (25.5%) and non-assigned reads (44.5%), with only ∼1% of the reads belonging to the class Amphibia. Average read depth for amphibian species per sample was 19,165 for the 12S vertebrate primer set and only 105 for the 16S amphibian primer set.

For the comparison of the primer sets, just the amphibian species detected are included. The 12S vertebrate primers detected six amphibian species where the 16S amphibian primers only detected five, missing *T. cristatus* which is a species of concern for these sites. *Lissotriton vulgaris* clearly dominates the reads and the two species of interest (*T. cristatus* and *P. fuscus*) are the least frequently detected species for both primer sets. See the Supplementary Material and Figure S1 for a breakdown of the proportion of reads for each amphibian as detected by each primer set.

Maps were constructed displaying the amphibian communities at each site as detected by the 12S vertebrate (Figure 2) and 16S amphibian (Figure 3) primer sets alongside the *Bsal* data, with 8 sites found to be *Bsal* positive and 19 to be *Bsal* negative. Figures 2 and 3 further show the deficiency of the 16S amphibian primer data as it contains ten sites with no amphibian reads where the 12S vertebrate primers detected amphibians. However, of the four sites where nothing was detected by the 12S vertebrate primers, the 16S amphibian primers detected species at three. At site 22 the 16S amphibian primers detected *L. vulgaris* but no sequence data was returned for this site by the 12S vertebrate primers. At site 16, the 16S amphibian primers detected *L. vulgaris* and at site 19 it also detected *R. temporaria* where the 12S vertebrate primers detected no amphibian species. However, all of these detections by the 16S amphibian primer set have a low number of reads, ranging between only 8 and 23 reads after filtering.

**Fig. 2.**
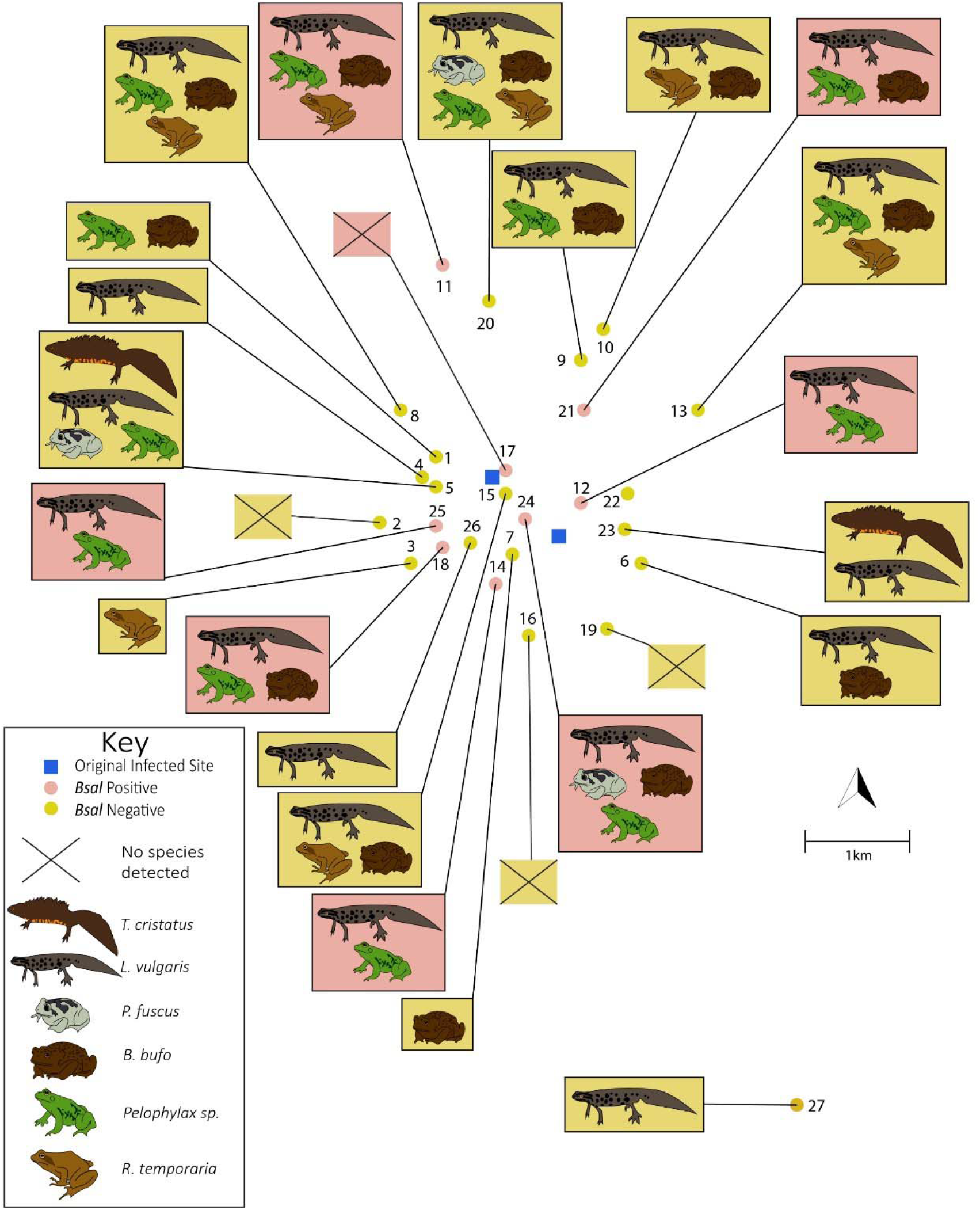
Map displaying the amphibian communities detected using the 12S vertebrate primer set at *Bsal* positive and negative sites (underlying map not shown since ponds are on private properties). The original two sites where *Bsal* was detected are included in blue squares. Site 22 failed so no results are shown for this site.

**Fig. 3.**
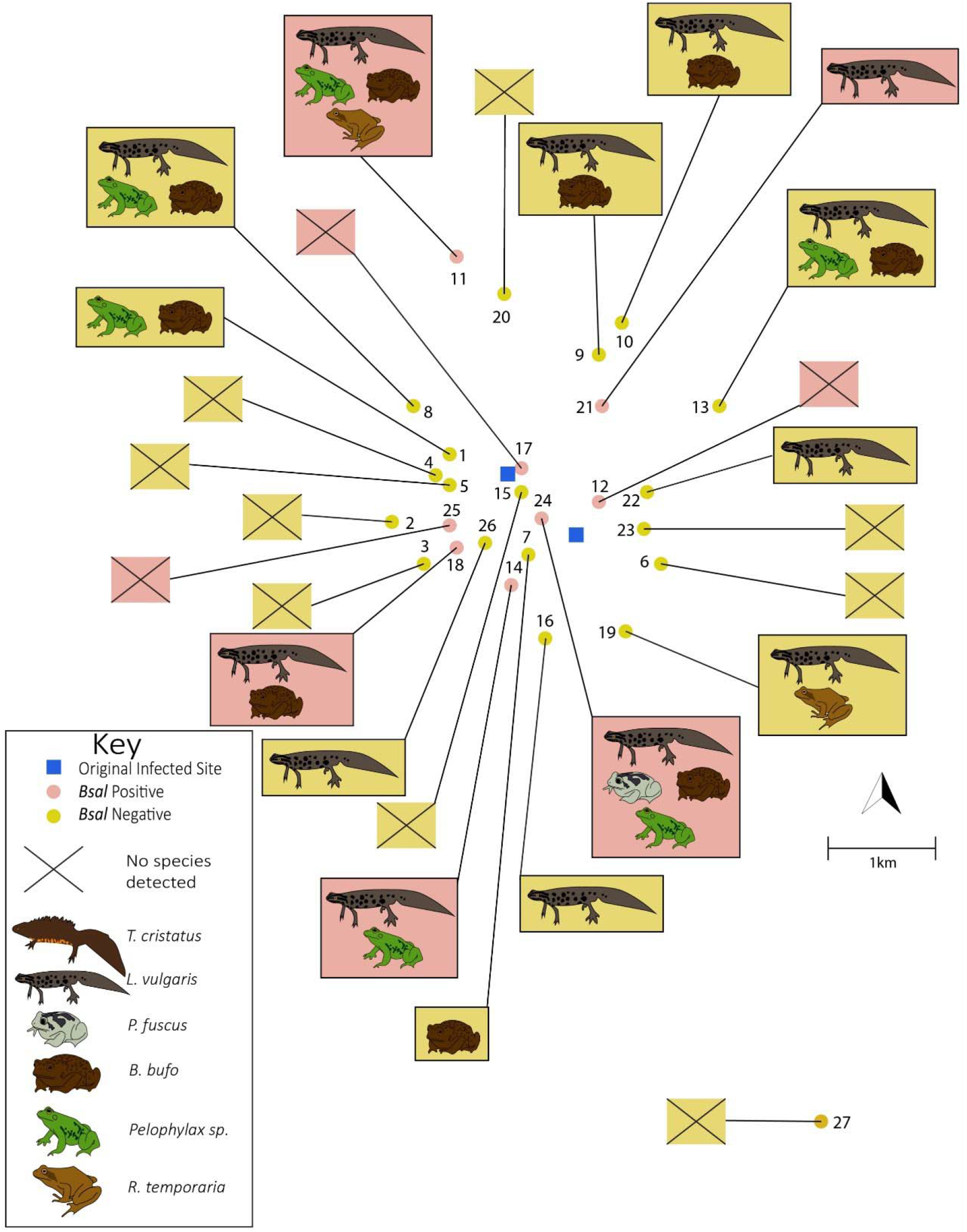
Map displaying the amphibian communities detected using the 16S amphibian primer set at *Bsal* positive and negative sites (underlying map not shown since ponds are on private properties). The original two sites where *Bsal* was detected are included in blue squares.

Overall, the 12S vertebrate primer set detected a generally higher species richness across sites than the 16S amphibian primer set with a median richness of two for the former and one for the latter. The 12S vertebrate primer set has been much more successful at detecting the species richness of the sites, reaching a plateau within the number of sites surveyed in this study (Figure 4). Furthermore, the 12S vertebrate primer data almost independently drives the combined data in Figure 4, making the contribution of the 16S amphibian primer data essentially redundant. Therefore, the analyses on associations between amphibian communities and *Bsal* presence was only conducted using the 12S vertebrate primer set.

**Fig. 4.**
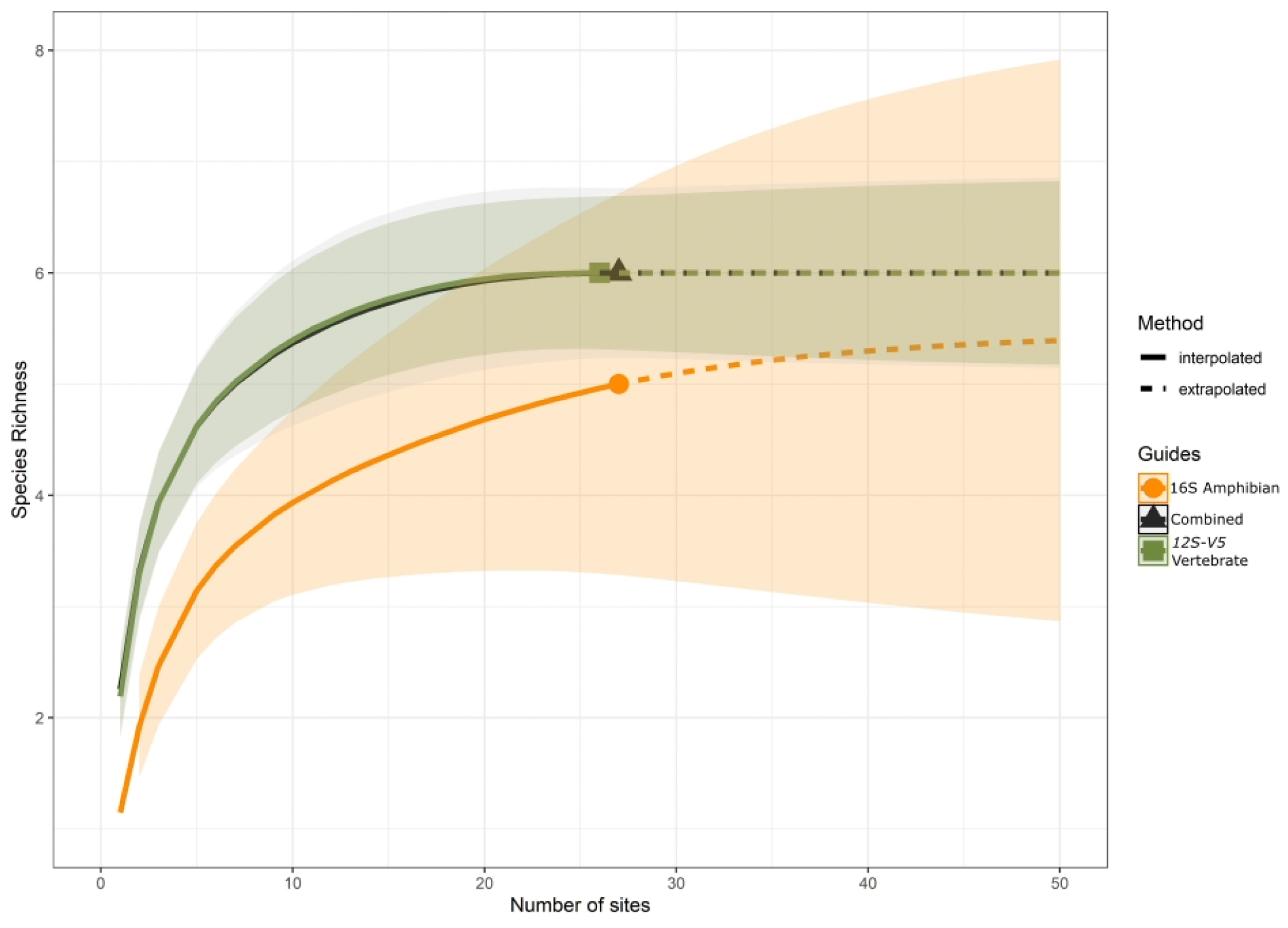
Species accumulation curve showing the interpolated species richness detected by each primer set and by both primer sets combined alongside the extrapolated richness to 50 sites. 12S vertebrate primer data has site 22 removed due to failure and therefore has 26 sites.

Sites were split into those which tested positive for *Bsal* and those which tested negative. The number of sites in each category is not equal, with 18 negative sites (site 22 is removed) and 8 positive sites. A PERMDISP test was conducted and the difference in distribution between *Bsal* positive and negative sites was found to be significantly different (F = 10.44, p = 0.005). This means that sites within the *Bsal* positive category have more intragroup similarity than those in the *Bsal* negative category. A dbRDA plot was created to explore the contribution of the *Bsal* infection presence to the species community composition observed. The dbRDA (Figure 5) was found to be significant (R^2^adj = 0.0569, p=0.0497). Of the maximum variance in the dataset (10.18%), the low adjusted R^2^ value suggests that the *Bsal* infection explains 5.7% of this total variance in amphibian community composition.

**Fig. 5.**
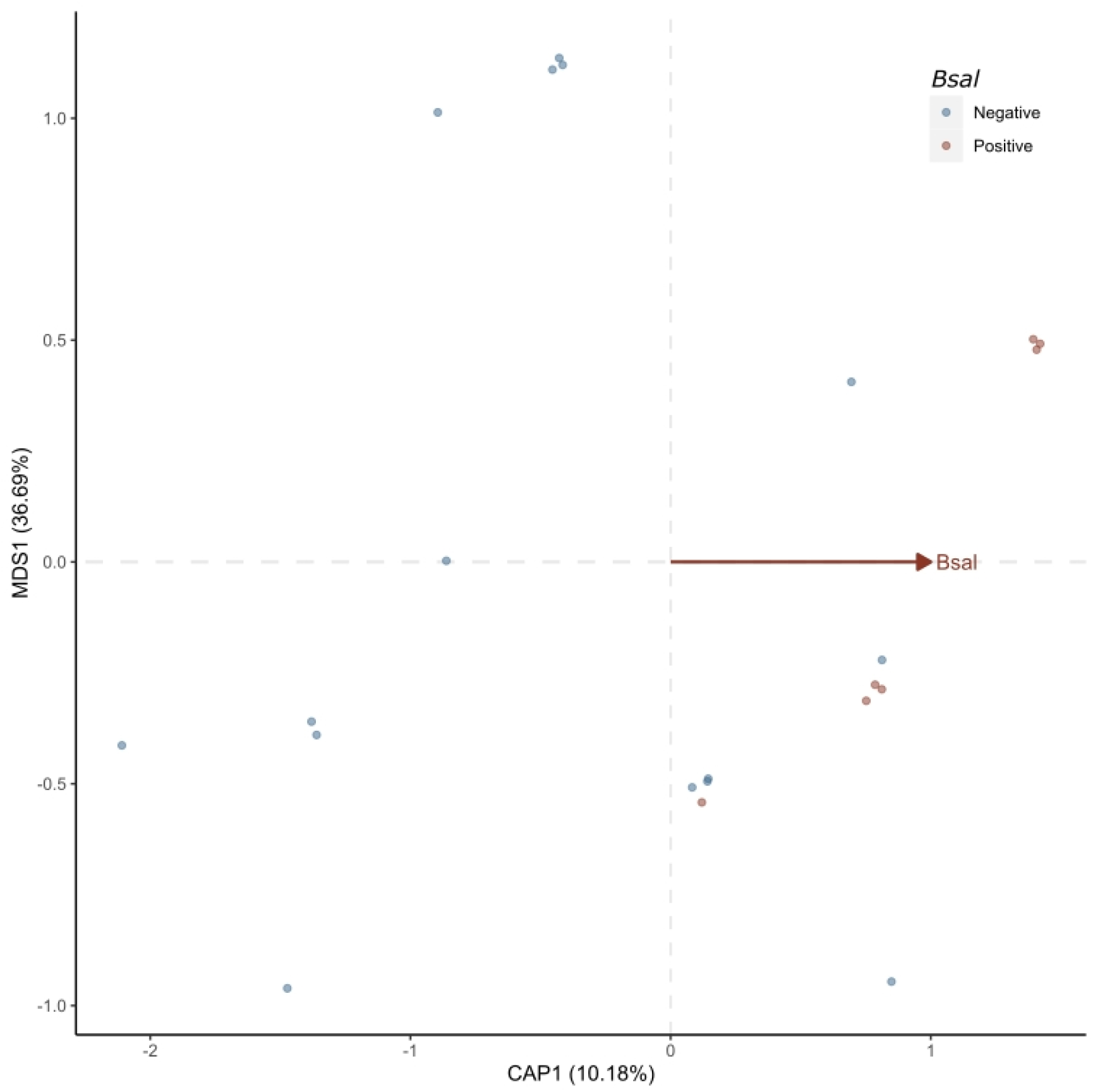
dbRDA plot showing the influence of *Bsal* on the differences in amphibian community composition as detected by the vertebrate 12S vertebrate primer set. Points are jittered for visibility.

## Discussion

This study has successfully detected the expected range of amphibian species at sites in Gelderland alongside a component of the wider vertebrate community present at these sites (Table S2). The species accumulation curve for the 12S vertebrate primer set (Figure 4) demonstrates that the overall sampling effort in this study was sufficient to detect all amphibian species expected in the study area. The *Bsal*-targeted qPCR demonstrates that *Bsal* has spread from the initial two sites to eight further sites without a clear path of spread as some of the closest sites to the original two sites remain uncontaminated (Figs. 2, 3). However, it should be noted that the chronological contamination of sites by *Bsal* is unknown as only two sites were sampled originally (RAVON, 2021).

Comparisons between the 16S amphibian and 12S vertebrate metabarcoding primer sets revealed that the former consistently failed to detect the full range of amphibian species detected by the latter at each site. The 16S amphibian primers did not detect *T. cristatus*, consistently detected a lower amphibian species richness at each site and contained <1% of the amphibian reads detected by the 12S vertebrate primers. The 16S amphibian primers have not yet been used in any published peer-reviewed studies except the original Bálint et al. (2018) study. It is likely that, as this primer set was developed for use within tropical biodiverse ecosystems and developed from anuran sequences from South America, it was not optimized for use in detecting European amphibian species. However, of greater concern is that this primer set leads to a large proportion of non-target amplification, with large quantities of Rotifera and Arthropoda being amplified (Table S1), which likely hinders its ability to optimally target amphibians (Collins *et al*. 2019). The 12S vertebrate primers did detect all of the expected amphibian species in the study area, with an average of 2.2 species per site, which is in line with a previous study in the Netherlands where one to five reproducing species were found per pond (Stumpel and van der Voet, 1998). However, this could possibly be an under-estimation of the species present at each site, with some of the rarer species being detected less frequently. For example, one of the main species of concern, the great crested newt *T. cristatus*, was not detected as often as perhaps it should have been given the known populations of *T. cristatus* in the area (A. Spitzen-van der Sluijs, pers. comm.). A previous study in the UK demonstrated that while these general 12S vertebrate primers are less sensitive than a species-specific qPCR assay for *T. cristatus*, it did provide similar proportions of detections when more conservative detection thresholds were applied (Harper et al., 2018). However, a more targeted approach (either with a species-specific qPCR assay or amphibian-focused 12S metabarcoding using the batra primers; Valentini et al., 2016) would be warranted in future studies. It is not currently known whether *T. cristatus* sub-populations have experienced any significant decline due to *Bsal* in this area. However, evidence from other studies have shown *T. cristatus* experiences a moderate mortality rate when exposed to *Bsal* and declines have been observed in wild German sub-populations (Bates et al., 2019; Lötters et al., 2020).

Having a record of the full amphibian communities present at sites of *Bsal* infection is valuable as the species at risk of infection and the potential vector species can then be determined. Furthermore, it is not yet known whether *Bsal* has an impact on the amphibian community as a whole. *Bd*-induced chytridiomycosis has caused huge declines in host species and the extirpation of many (Lips et al., 2006; Scheele et al., 2019) which then leaves an empty niche which can be utilized by other species (Hirzel and Le Lay, 2008). A similar outcome, but on a smaller scale, could occur at sites of *Bsal* infection. If a *Bsal*-tolerant species fills this niche, it could lead to an acceleration of *Bsal* transmission but if a *Bsal*-resistant species fills the niche, it can act to slow the spread of the fungus (Holt and Pickering, 1985; Brannelly et al., 2021). In this study, *L. vulgaris* can be considered a tolerant species (Bates et al., 2019) and anurans are considered resistant (Martel et al., 2013). The potential trend between *Bsal* presence and the community composition of amphibians was investigated using a dbRDA, which concluded that the *Bsal* infection explained only a small proportion of the variance in the amphibian community composition detected here. Well known drivers such as habitat and microhabitat availability suitable for amphibian breeding activities are likely to be much more influential on amphibian community composition than *Bsal* here (Werner et al., 2007; Vági et al., 2013; Konowalik et al., 2020). However, the significant difference in the intra-group variability, with *Bsal* positive sites containing amphibian communities which are more similar, could potentially show a homogenization of communities due to *Bsal* or that these similar communities are more likely to be successfully infected with the fungus (Smith et al., 2009). This is clearly an avenue which requires further investigations.

There are a number of reasons why, with further data, *Bsal* may yet be found to be an important driver of amphibian community composition. Firstly, there is the caveat that the eDNA metabarcoding approach used here may be under-estimating the amphibian community in the area (see discussion above). Secondly, this study contained relatively few sites and only one sampling occasion. Larger spatial and temporal resolution would provide the volume of data needed to accurately reveal the relationship between *Bsal* and amphibian community compositions (Beentjes et al., 2019). As most amphibians in this study vacate the water outside of spring and early summer, sampling times are still advised within the amphibian breeding season to ensure sufficient eDNA present in the water but sampling over many years could track changes in communities as they occur. Detection of amphibian communities across *Bsal* positive sites in the Netherlands, ideally also including *Bsal* positive sites in Germany, Belgium and Spain, would provide the most comprehensive view of which species are exposed to the pathogen in its invasive range. It would also allow for more in-depth analyses into the dynamics of the amphibian communities within *Bsal* positive areas and potentially enable the construction of a chronosequence in the absence of long-term studies to provide further insight into community development over time.

Abundance data would also have added another dimension to this community analysis and perhaps revealed differences in the populations of species at *Bsal* positive and negative sites. It could also have revealed the status of *T. cristatus* populations, with the opportunity to track any declines if more sampling sessions are conducted in the future. One shortcoming of eDNA is the difficulty in consistently determining the abundance of species within a site or study area. Several studies have shown that read count is positively correlated with abundance or biomass, with fish being by far the most studied taxa (e.g. Takahara et al., 2012; Evans et al., 2016; Lacoursière-Roussel et al., 2016; Wilcox et al., 2016) and amphibians being the subject of only a few studies (Evans et al., 2016; Li et al., 2021). However, a recent meta-analysis found that fish are currently the only vertebrates where it can be reliably concluded that this positive correlation with abundance or biomass exists (Carvalho et al., 2021).

As expected, the rarest species detected in this study were those protected under the European Habitats Directive (Council Directive 1992/43/EEC): *Triturus cristatus* (Annex II and IV) and *Pelobates fuscus* (Annex IV). *Lissotriton vulgaris* dominates the read count in this study and is present at most sites which was to be expected as it is a common species and syntopic with *T. cristatus*. The proportion of *Bsal* positive and negative sites hosting each amphibian species and waterbirds as a group were examined but this study did not contain enough *Bsal* positive sites for further meaningful statistical analysis (data not shown). Even so, this work can be considered a preliminary study providing the groundwork for future research and provides several lines of future investigation. There are three main important groups identified within this study which require further research to determine their exact level of influence on *Bsal* transmission and potency. The first of these groups is represented here by the tolerant urodelan species *L. vulgaris.* The little research done on *L. vulgaris* and *Bsal* suggests that the species is not as prone to developing lethally high loads of the fungus as *T. cristatus* and is more likely to fully recover (Bates et al., 2019). Therefore, *L. vulgaris* could help drive *T. cristatus* decline by acting as a reservoir, constantly providing a source of new infection for *T. cristatus* populations (Brannelly et al., 2021; Lötters et al., 2020). Furthermore, this species is widespread throughout the study area, and the two species are highly associated, sharing certain prey species and hibernacula (Roşca et al., 2013; Dervo et al., 2018). *Bsal* is likely to spread between them despite the difference in their aquatic spatial niches (Dolmen and Koksvik, 1983; Covaciu-Marcov et al., 2010) as spores are released into the environment (Stegen et al., 2017). Despite the capacity for a higher infection load (EFSA Panel on Animal Health and Welfare, 2018), it is unknown if *L. vulgaris* is acting as an effective vector for *Bsal* as it generally travels much shorter distances than the anuran species detected here (Kovar et al., 2009). However, *L. vulgaris* can reach high densities (Bell and Lawton, 1975) so it could potentially be a key component of *Bsal* transmission and persistence in the Netherlands, acting in much the same way as the introduced *Triturus anatolicus* population is in the newly discovered *Bsal* outbreak in Spain (Martel et al., 2020).

The second important group identified in this study is the anurans which are passive carriers of *Bsal* and cover longer distances during migration than *L. vulgaris*, making them potential vectors for *Bsal* within the study area. *Bufo bufo, R. temporaria* and *Pelophylax spp.* are all capable of travelling several kilometers overland, mostly for the purposes of breeding (Juszcyk, 1951; van Gelder et al., 1986; Tunner, 1992; Kovar et al., 2009). Of these anurans, only *Pelophylax spp.* was present at all amphibian-occupied *Bsal* positive sites (Figure 2). Since these distances are mostly travelled during the spring when individuals migrate from winter hibernacula to breeding ponds and between ponds (Juszcyk, 1951; Tunner, 1992; Kovar et al., 2009), this hypothesis of anuran transmission of *Bsal* between sites relies upon the persistence of the fungus over winter in hibernacula and on the skin of vector or host species. Although a direct study testing overwintering amphibians for the presence of *Bsal* has yet to be conducted, *Bsal* has been shown to thrive at low temperatures on terrestrial species (*S. salamandra*; Martel et al., 2013) and therefore is hypothesized by experts to be capable of spreading through overwintering and subsequent migration (EFSA Panel on Animal Health and Welfare, 2018). However, anurans are known to have lower infection loads compared to urodelans and therefore validation that *Bsal* can indeed persist on overwintering anurans is needed.

The final important group to consider is the potential long-distance vectors. Although the contribution of amphibian-amphibian transmission cannot be discounted, barriers to amphibian movement, such as roads and buildings, are very common in the study area making long-distance travel less likely. Stegen et al. (2017) have shown that waterbirds could be effective long-distance vectors of *Bsal* and the presence of waterbirds at *Bsal* positive sites in this study (Table S2) supports the need for further research to quantify their impact. Although difficult to quantify, perhaps the most likely culprit for long-distance *Bsal* transmission could be humans. Many of the water bodies sampled in this study are on small plots of private land and therefore are highly likely to come into frequent contact with humans. The impact of interactions with the sites by RAVON when sampling or visiting was minimized by thorough decontamination of the equipment using protocols deemed to be sufficient to prevent spread (Thomas et al., 2019; RAVON, 2020).

In conclusion, eDNA metabarcoding is a suitable method for detecting amphibians in the Netherlands and the full range of amphibian species expected in the study area were detected. The dbRDA determined that *Bsal* was significant as an explanatory variable for amphibian community composition but further investigations incorporating variables such as suitable habitat and microhabitat availability are clearly warranted. Despite the small dataset, several lines of enquiry were identified and discussed for future studies. *Lissotrition vulgaris* has the potential to exacerbate the impact of *Bsal* on *T. cristatus* populations by acting as a reservoir. If *Bsal* can endure cold winters on their skin, anurans could be effective vectors for *Bsal* between ponds during the spring travel from hibernacula to breeding ponds. Potentially more effective long-distance vectors of *Bsal* are waterbirds and humans which are the most likely culprit for the transmission between widely-distributed ponds and those with barriers to amphibian movement. Once the transmission of *Bsal* is understood, new sites of contamination could be predicted and measures can be taken to slow the spread and prevent further mass mortality events.

## Acknowledgements

We thank the landowners for permission to sample ponds on their property. We thank the members of the Molecular Ecology Group in the University of Salford for their advice on laboratory protocols and bioinformatics.

