## Supplementary Material for "Using Environmental DNA to Reconstruct Amphibian Communities at Sites Infected with *Batrachochytrium salamandrivorans* in the Netherlands"

**Library Preparation, Sequencing and Bioinformatics**

RATIO Agencourt AMPure XP (Beckman Coulter) was used to perform a left-sided size selection with a bead ratio of 1:1.5 for the *12S-V5* vertebrate primers library and 1:0.9 for the 16S amphibian primers library. The KAPA HyperPrep kit (KapaBiosystems) was used to add Dual-Index adapter 4 to each library. The NEBNext qPCR quantification kit (Biolabs) was then used to quantify each library by qPCR in order to dilute correctly to a concentration of 9pM with 1% PhiX in preparation for sequencing. An Illumina MiSeq Reagent v2 2x150-bp nano kit was then used to sequence each library on separate sequencing runs.

The bioinformatic analysis was conducted using the OBITools metabarcoding package (Boyer *et al.,* 2016) followed by a direct blast against the GenBank nucleotide database (NCBI) to assign taxonomy. For the taxonomic assignment, MOTUs required at least 98% identity to be assigned at species level, retained more than five reads and all other assignments were filtered out in subsequent steps (Sales *et al.,* 2020).

The results were filtered using an adapted version of the method used in Broadhurst *et al.* (2021) with read counts at each stage recorded in Table S1. First, reads were removed proportional to the number of positive control reads found in samples other than the positive control to account for tag jumping (Schnell *et al.,* 2015). Then the maximum number of reads within the negative control samples were removed from all read counts. Reads from domestic animals and humans were removed. MOTUs assigned at species level and with more than five reads were kept to produce a file containing the vertebrate community. Subsequently, the amphibian community was isolated in a separate file. For the 16S amphibian primers, the same steps were followed but no vertebrate community file was created. *Pelophylax lessonae* was replaced with *Pelophylax spp*. as the mitochondrial DNA of *P. lessonae* and the hybrid *Pelophylax kl. esculentus* are indistinguishable using eDNA methods (Holsbeek and Jooris, 2010), although the hybrid is more common.

**Table S1.** Read counts for each primer set at each stage of the post-bioinformatic filtering process.

|  | 12S Vertebrate | 16S Amphibian |
| --- | --- | --- |
| Total reads before filtering criteria | 1370905 | 542354 |
| After accounting for tag-switching | 1368503 | 542354 |
| After removing positive control | 1366027 | 530744 |
| After removing negative control | 1365335 | 521921 |
| After removing all human reads | 998859 | 521423 |
| After removing domestic animals (Bos, Canis, Felis, Ovis and Sus) | 963799 | 521423 |
| MOTUs with minimum identity >0.98 and less than 5 reads removed | 945831 | 99420 |
| After removing all non-amphibian reads | 498302 | 2848 |
| Tag-switching proportion | 0.0004 | 0 |

The vertebrate community produced by the vertebrate primers was extracted just before removing all non-amphibian reads in the filtering process and is shown in Table S2. All three replicates for site 22 failed and it was subsequently removed from the data. Thirteen species of birds were detected including five waterbirds: mallard (*Anas platyrhynchos*), grey heron (*Ardea cinerea*), Eurasian coot (*Fulica atra*), white stork (*Ciconia Ciconia*) and great cormorant (*Phalacrocorax carbo*). One of the bird species detected was the Siberian accentor (*Prunella montanella*), a rare migrant to Western Europe. The sequence has 234 reads at one site and was a 100% match with *P. montanella* and *Prunella fulvescens* but the former was deemed more feasible. Eight mammal species were detected with most species occurring at just one site except the beech marten (*Martes foina*), wood mouse (*Apodemus sylvaticus*) and roe deer (*Capreolus capreolus*) which occur at two sites. Four fish species were detected but only the asp (*Leuciscus aspius*) is native in the wild in Europe (but introduced to the Netherlands) and therefore the other three species are assumed to have been introduced into sites as decorative pond species. The results from RAVON and SPYGEN showed that eight of the 27 sites tested positive for *Bsal* and these are also displayed in red in Table S2.

**Table S2.** Table with green boxes indicating detection of a species by the *12S vertebrate* primer set. *Bsal* positive sites are indicated in red and site 22 is shaded grey due to failure. Species in red are non-native in the Netherlands.

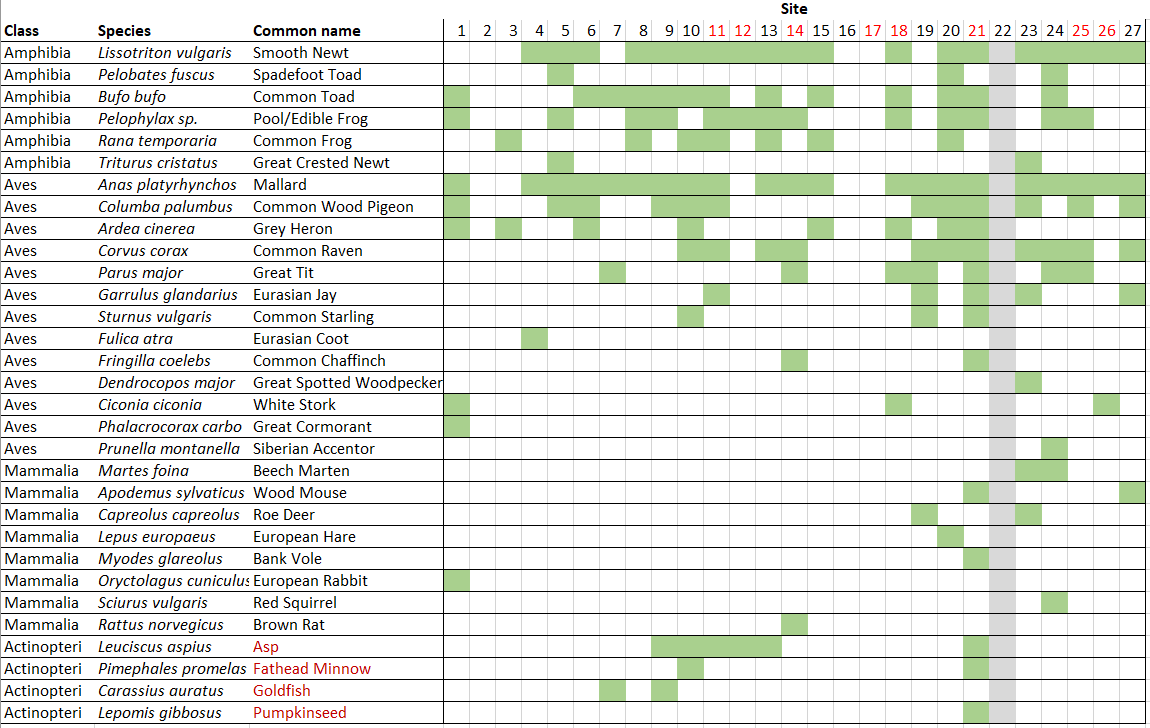

The proportion of reads which constitutes each species at each site for both primer sets is shown in Figure S1. There are some similarities in the proportion of each species detected by both primer sets, with a few sites showing almost the same proportion and composition for both primer sets such as 1 and 24.

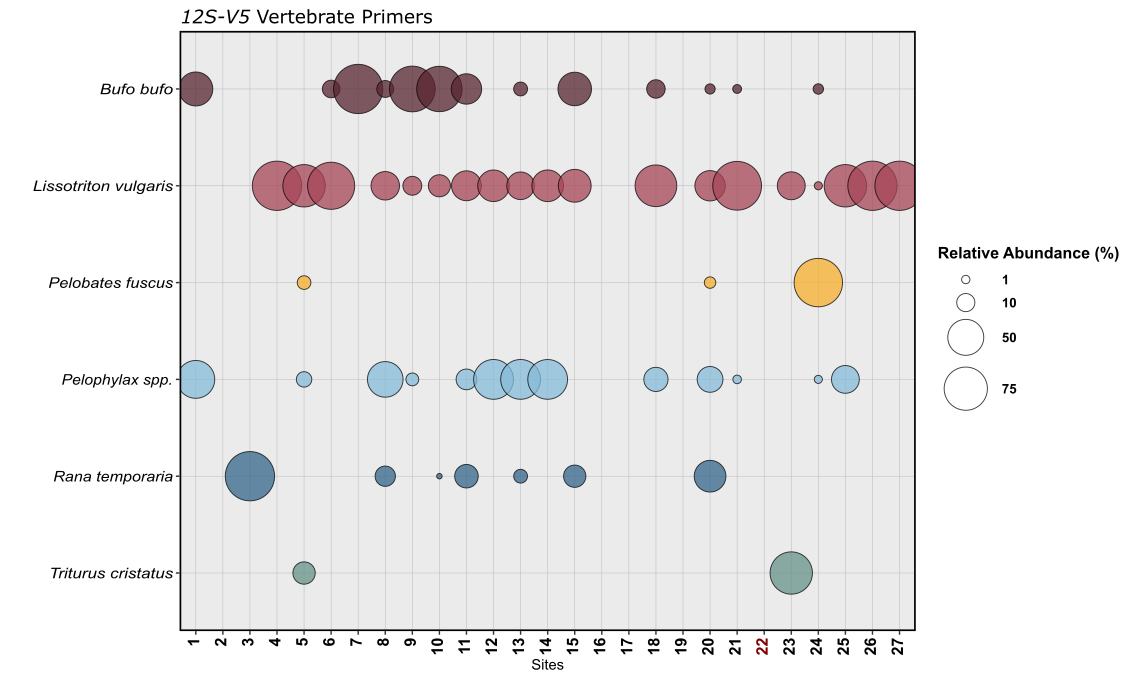

A

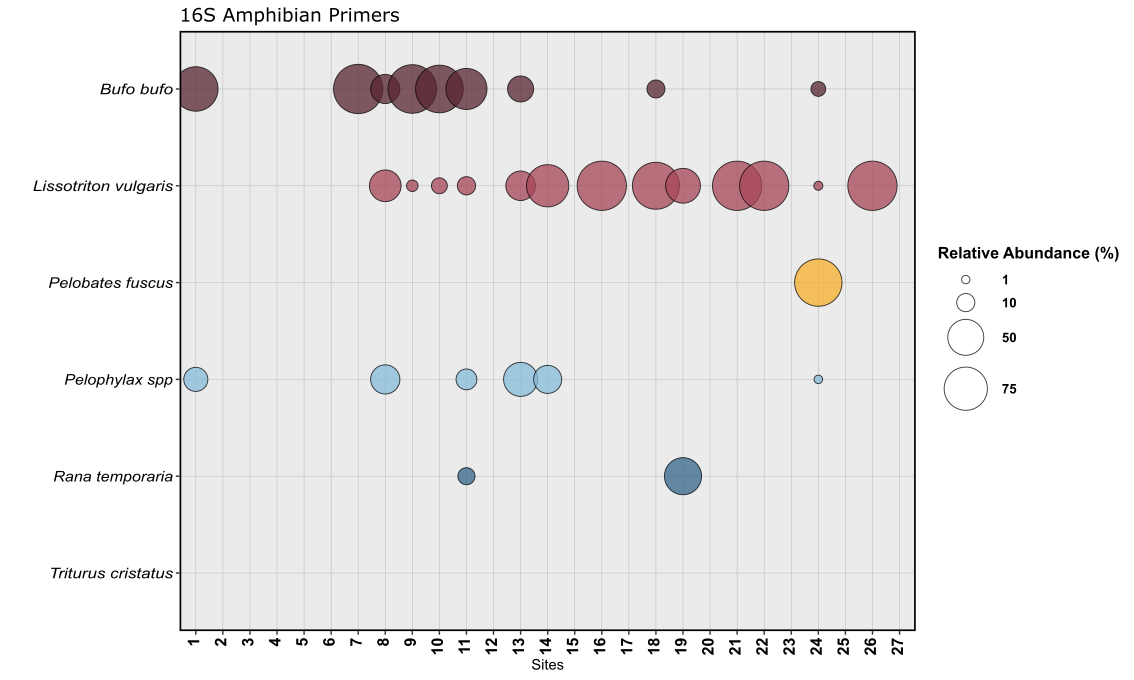
**Figure S1.** Relative abundance of the reads for each species at each site detected using the (A) *12S vertebrate* primer set and (B) *16S amphibian* primer set. Site 22 (red) failed for the vertebrate primer set and was subsequently removed from the data.

B
